# The heterogeneous nature of the Coronavirus receptor, angiotensin-converting enzyme 2 (ACE2) in differentiating airway epithelia

**DOI:** 10.1101/2020.07.09.190074

**Authors:** Vincent J. Manna, Salvatore J. Caradonna

## Abstract

Coronavirus Disease 2019 (COVID-19) is transmitted through respiratory droplets containing Severe Acute Respiratory Syndrome Coronavirus 2 (SARS-CoV-2) particles. Once inhaled, SARS-CoV-2 particles gain entry into respiratory ciliated cells by interacting with angiotensin converting enzyme 2 (ACE2). It is known that ACE2 functions within the renin-angiotensin system to regulate blood pressure, fluid homeostasis and inflammation. However, it is largely unknown what roles ACE2 has in ciliated cells of the airway. Therefore, understanding the function and nature of ACE2 within airway tissue has become an essential element in combatting the COVID-19 pandemic. Airway mucociliary tissue was generated *in-vitro* using primary human nasal epithelial cells isolated from nasal turbinates of donors and the air-liquid interface (ALI) model of differentiation. Using ALI tissue we cloned transcripts for three distinct variants of ACE2, one of which encodes the full-length ACE2 protein, the other two transcripts are truncated isoforms that had only been predicted to exist via sequence analysis software. We demonstrate that all three isoforms have the capacity to be glycosylated, a known modification of full-length ACE2. Immunofluorescence microscopy of individual ACE2 isoform transfected cells reveals distinct localization of variant 1 relative to X1 and X2. Double staining immunohistochemistry of ALI tissue using antibodies to either the N-term or C-term region of ACE2 revealed distinct and overlapping signals in the apical cytosol of ciliated cells. Most notably only the ACE2 C-term antibody displayed plasma-membrane localization in ciliated cells. We also observed a decrease in the total amount of ACE2 in ALI tissue derived from a 33 year-old male donor when compared to a 34 year-old female donor, thus there may be variation in the abundance of ACE2 protein in the airway among the population. Together, our data begins to highlight the dynamic status of the ACE2 protein in airway mucociliary tissue and we propose multiple ACE2 parameters that may impact an individual’s susceptibility to SARS-CoV-2. These parameters include the balance of cytosolic versus membrane bound ACE2, isoform expression levels, maintenance of post-translational modifications and the impact of genetic, environmental and lifestyle factors on these processes.

## INTRODUCTION

The identification of the ACE2 protein as the receptor for SARS-CoV-2 infection leads to the tenet that the efficiency of infection is directly related to the amount and nature of ACE2 available for binding [1]. The main route of transmission of SARS-CoV-2 is through respiratory droplets, once inhaled the viral particles bind to and infect proximal airway cells using a viral encoded spike protein (S) that interacts with ACE2. ACE2 is an integral membrane protein attached to the outer surface of cell membranes [2, 3].

In addition to providing a receptor for SARS-CoV-2 infection, ACE2 plays a very seminal role in the body. As a counterpart to ACE, ACE2 has emerged as an important co-regulator of the renin-angiotensin system (RAS) [4]. The primary role of ACE is to hydrolyze angiotensin I (Ang I) to a potent vasoconstrictor, angiotensin II (Ang II). ACE2 functions to cleave Ang II to Ang (1-7), a peptide with opposing functions to Ang II that acts through its own receptor, MAS [5-7]. Ang II was originally considered as a short-acting, vasoactive hormone. Research over the years now presents Ang II as a peptide hormone that impinges on multiple signaling pathways. It can produce aberrant effects leading to cardiovascular disease with pathologies that include cardiac hypertrophy, endothelial dysfunction, fibrosis and inflammation [8]. Accumulating evidence indicates that Ang-(1-7) can reduce the levels of pro-inflammatory cytokines such as TNF-alpha and IL-6 [9]. In addition, data from ACE2 knockout mice suggests that ACE2 plays a protective role in acute lung injury [10]. This evidence indicates that ACE2 has a role in moderating inflammatory responses. In addition ACE2 is also known to neutralize other vasoactive peptides such as bradykinin [4]. Additional apparent roles for ACE2 include stabilization of cell adhesion through binding to integrin subunits and binding interactions with calmodulin which is thought to stabilize ACE2 at the cell surface [11, 12].

Considering ACE2 function in the airway, Jia et al. demonstrated that airway tissue releases a soluble form of ACE2 (sACE2) from the cell surface into the mucosal layer, shedding occurs by proteolytic cleavage of the ectodomain from the membrane. Recently it has been shown that sACE2 can partially block SARS-CoV-2 infection of ACE2 expressing cells [13, 14]. The release of sACE2 from epithelia is both constitutive and inducible [15] [16]. Inducible release is instigated by a number of factors including inflammatory stimuli. Calmodulin inhibitors increase sACE2 production, substantiating an association between these two functions [12]. Earlier research on SARS-CoV has demonstrated that this virus can induce shedding of sACE2 protein from cells [17]. Furthermore, the SARS-CoV spike protein, independent of the virus, induced shedding of sACE2 from cells. Work by Kuba et al. showed that the SARS-CoV spike protein, injected into mice worsens acute lung failure *in vivo* that could be attenuated by blocking the renin-angiotensin pathway [18]. Conclusions drawn from these studies indicate that disruption of ACE2 allows elevated levels of Ang II to produce the pathological effects, described above.

In this study we utilized the air-liquid interface model of differentiation to generate airway mucociliary tissue using human nasal epithelial cells (HNECs) isolated from nasal turbinates of volunteer donors. Using cDNA generated from ALI tissue we cloned out transcripts for three isoforms of ACE2, each differing at the carboxy-terminus of the protein. Through transfection and PNGaseF treatment we establish that all three ACE2 isoforms are able to be glycosylated. These isoforms appear to display distinct morphology when introduced individually into cells. Fluorescent immunohistochemistry also indicates heterogeneity of ACE2 within ALI tissue. We observed ACE2 located both in the basal cell compartment and the ciliated cell compartment, but minimal signal in the central region of tissue. The apical cytosol of ciliated cells contained a mixture of cytosolic and membrane-bound ACE2. When we compared ACE2 immunohistochemistry patterns between ALI tissues derived from different donors we observed variation in overall amount and intensity, indicating potential variation in ACE2 expression between individuals. Together, our studies highlight the heterogeneity of ACE2 status in the airway within an individual, as well as variation in these parameters between individuals.

## RESULTS

### Identification of predicted ACE2 isoforms in ALI Tissue

Genbank reports two variants of human ACE2, v.1 and v.2 (NM_001371415 and NM_021804) which have identical coding sequence but differ in the 5’ UTR. There are three predicted isoforms of ACE2, v.X1, v.X2 and v.X3 (XM_011545549, XM_011545551 and XM_011545552). We designed cloning primers specific to the open-reading frames of the individual isoforms, then generated cDNA from fully-differentiated HNEC ALI tissue and performed PCR reactions with our isoform-specific primer pairs. We sequenced and cloned three distinct isoforms of ACE2 (v.1/v.2, v.X1 and v.X2) into expression vectors. These isoforms code for proteins that vary at their extreme carboxy-terminus (Figure 1). We did not identify the shorter v.X3 isoform.

**Figure 1:**
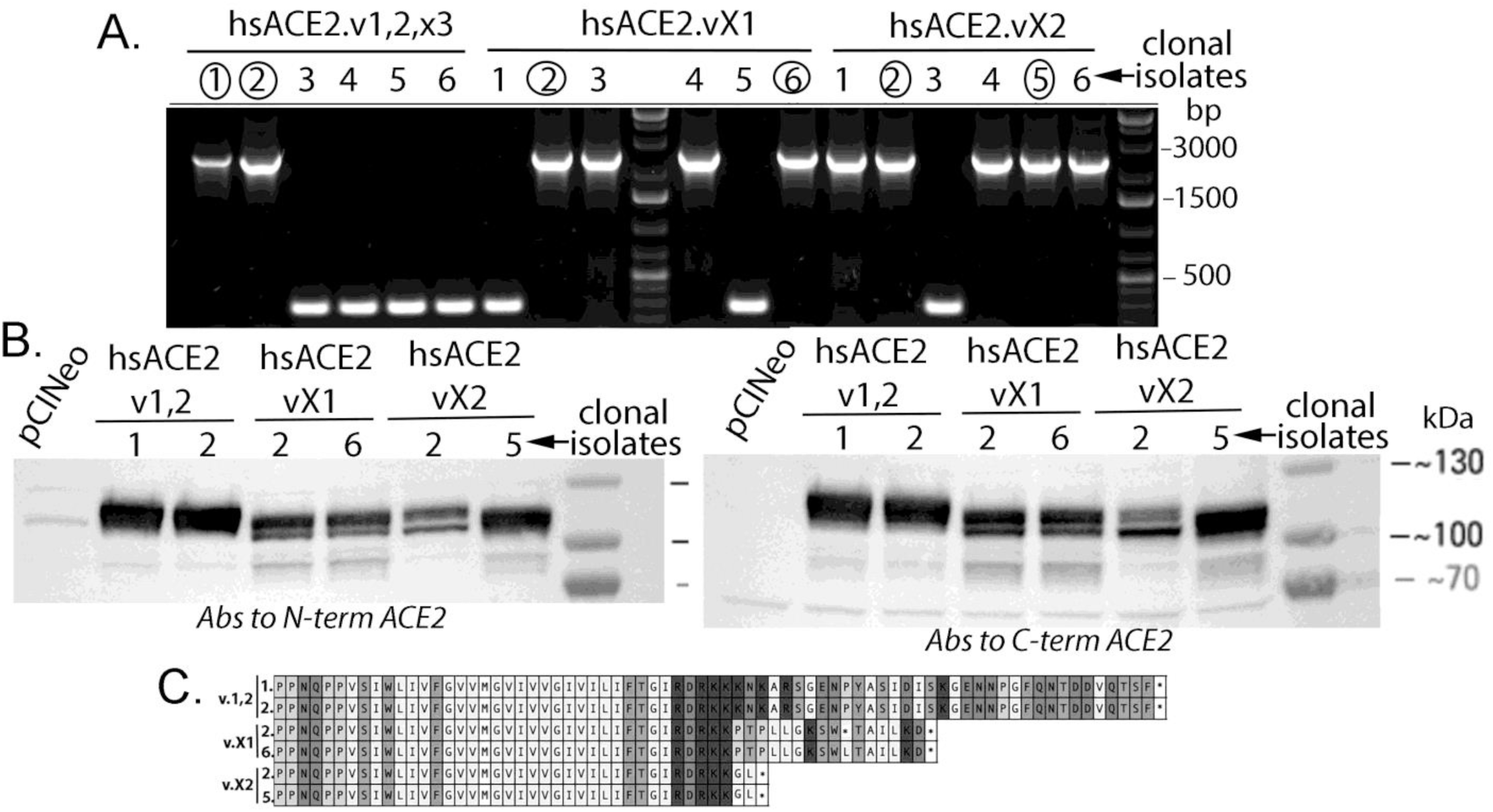
Identification of three isoforms of human ACE2. A.) Colony PCR analysis of recombinant clones derived from 3’ primers, specific for variants 1, 2 and X3; X1 and; X2 revealed the presence of distinct ACE2 isoforms. Two clonal isolates of each variant were analyzed further. B.) Transfection into cells of the cloned isoforms and western blot analysis reveal that they all express protein of approximately 120 kDa. Antibodies generated to either the N-terminus (ProSci 3227) or C-terminus (Proteintech 66699) of ACE2 give essentially identical results when used in western blot analysis. C.) DNA sequence analysis reveals that the three variants differ at the C-terminus in both sequence length as well as amino acid sequence. Amino acid sequence shown represents residues 733 to 805 of the full length ACE2 (variant 1,2).

The expression vector PCINeo, containing the individual isoforms was transfected into HEK293 cells and subsequent ACE2 immunoblotting of cell extracts revealed bands varying in sizes of about 120 kDa. When the cell extracts of 293 transfects were treated with PNGaseF to remove any glycosylation modifications the bands observed by immunoblot shifted to ∼100kDa, indicating all three isoforms are capable of modification via glycosylation (figure 2).

**Figure 2:**
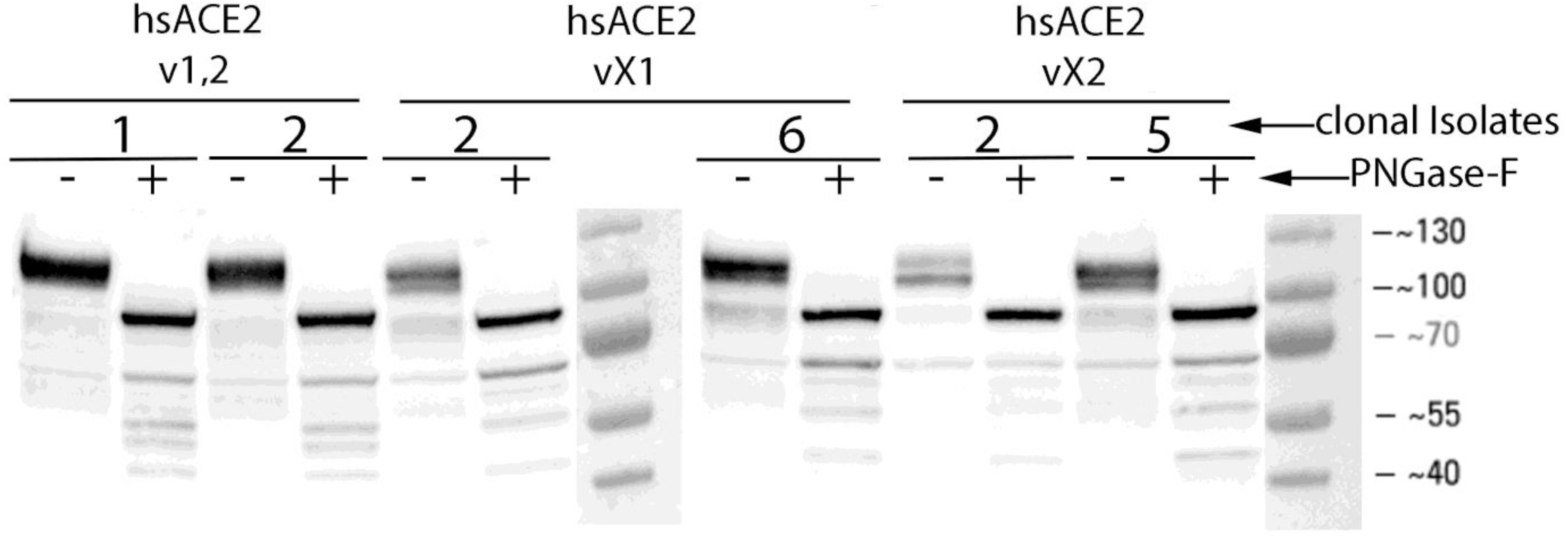
Demonstration of glycosylated modification of the three isoforms of ACE2. Clonal isolates identified (two each) as the three isoforms were subjected to digestion with PNGaseF. As illustrated PNGaseF treatment results in a shift in molecular weight from about 120 kilodaltons to about 100 kilodaltons.

### Immunofluorescent microscopy of the individual isoforms of ACE2 reveal distinct morphologies

Individual isoforms of ACE2 were transfected into HEK293 cells and visualized by immunofluorescence microscopy. As seen in figure 3 the isoforms appear to distribute differently within the cell. Full length ACE2 (variant 1,2) localizes to the plasma membrane distributed largely as foci. In contrast variants vX1 and vX2 appear cytosolic with an intense perinuclear distribution. This data demonstrates the intrinsic ability of these isoforms to localize to different areas of the cell.

**Figure 3:**
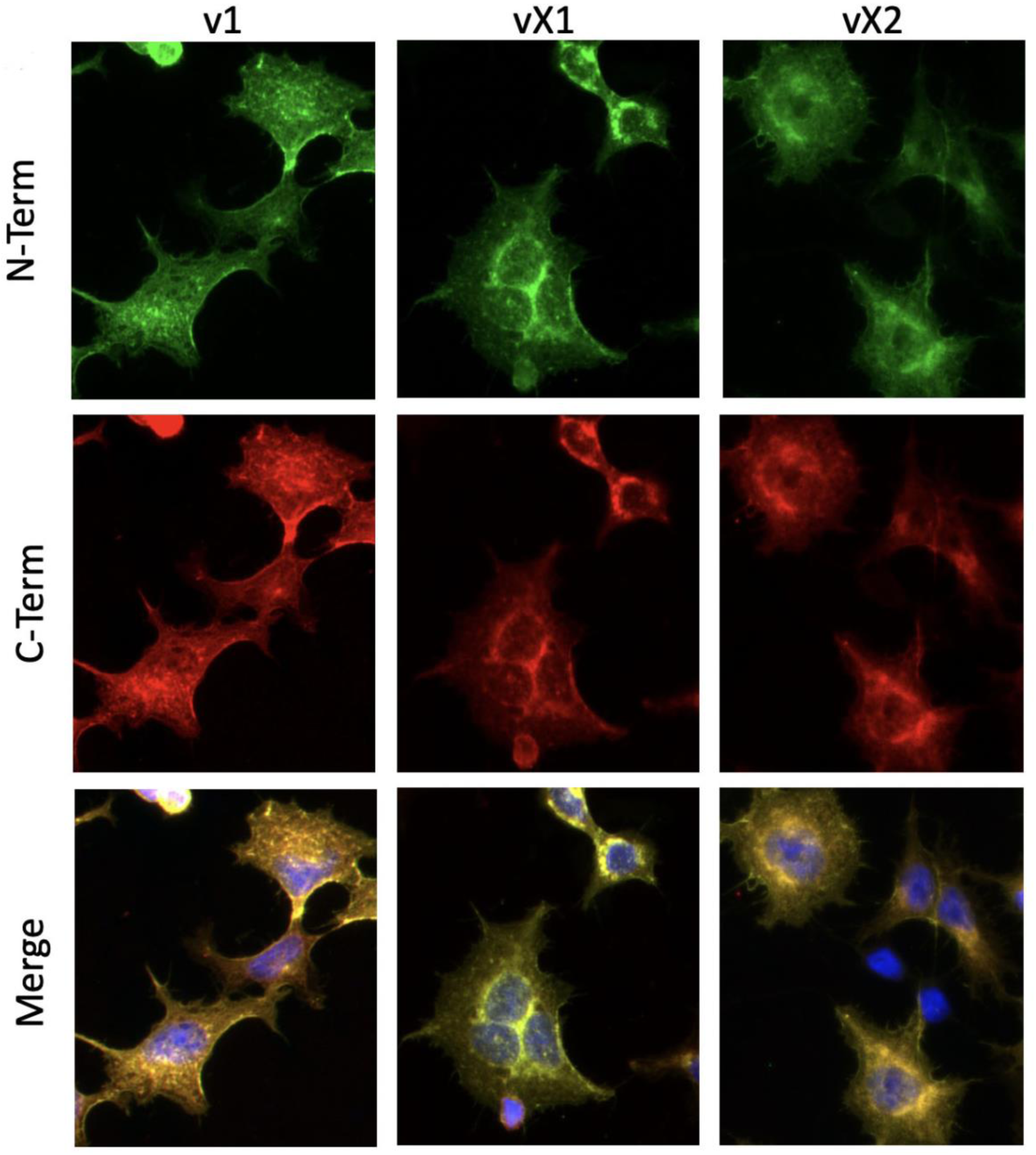
Immunofluorescence microscopy of individual isoforms of ACE2. HEK293 cells were transfected with each of the isoforms of ACE2 and then processed for immunofluorescent microscopy. As can be seen in the figure above, there are distinct localizations of the isoforms. Green; N-terminus antibodies (Proteintech 66699, 1:500 dilution). Red; C-terminus antibodies (ProSci 3227, 1:250 dilution).

### ACE2 expression in HNEC progenitor cells and differentiating ALI tissue

HNEC cells were analyzed by immunoprecipitation and western blot to elucidate ACE2 expression in these cells and during the differentiation phase of an ALI culture. 500 µg of progenitor cell protein extract, as well as 500 µg each of day 8 and day 18 ALI growth were subjected to immunoprecipitation and western blot analysis. Progenitor cells growing in submerged conditions (i.e. media covering the cells) do not show any ACE2 protein at the limits of detection in our system. This is consistent with our observations of ACE2 expression at the RNA level. We do not find ACE2 transcript from analysis of several cDNA libraries constructed from progenitor HNECs that were grown under submerged conditions. This work verifies earlier findings indicating little to no expression of ACE2 in airway basal cells [15]. However, when HNEC progenitor cells are exposed to an air-liquid interface for 8 days ACE2 protein becomes apparent. This data indicates that HNEC cells, when maintained under progenitor conditions, express minimal to no ACE2 protein. Analysis of a shorter time-course of ALI growth reveals that ACE2 expression begins at least at day 2 and increases thereafter (figure 4B).

**Figure 4:**
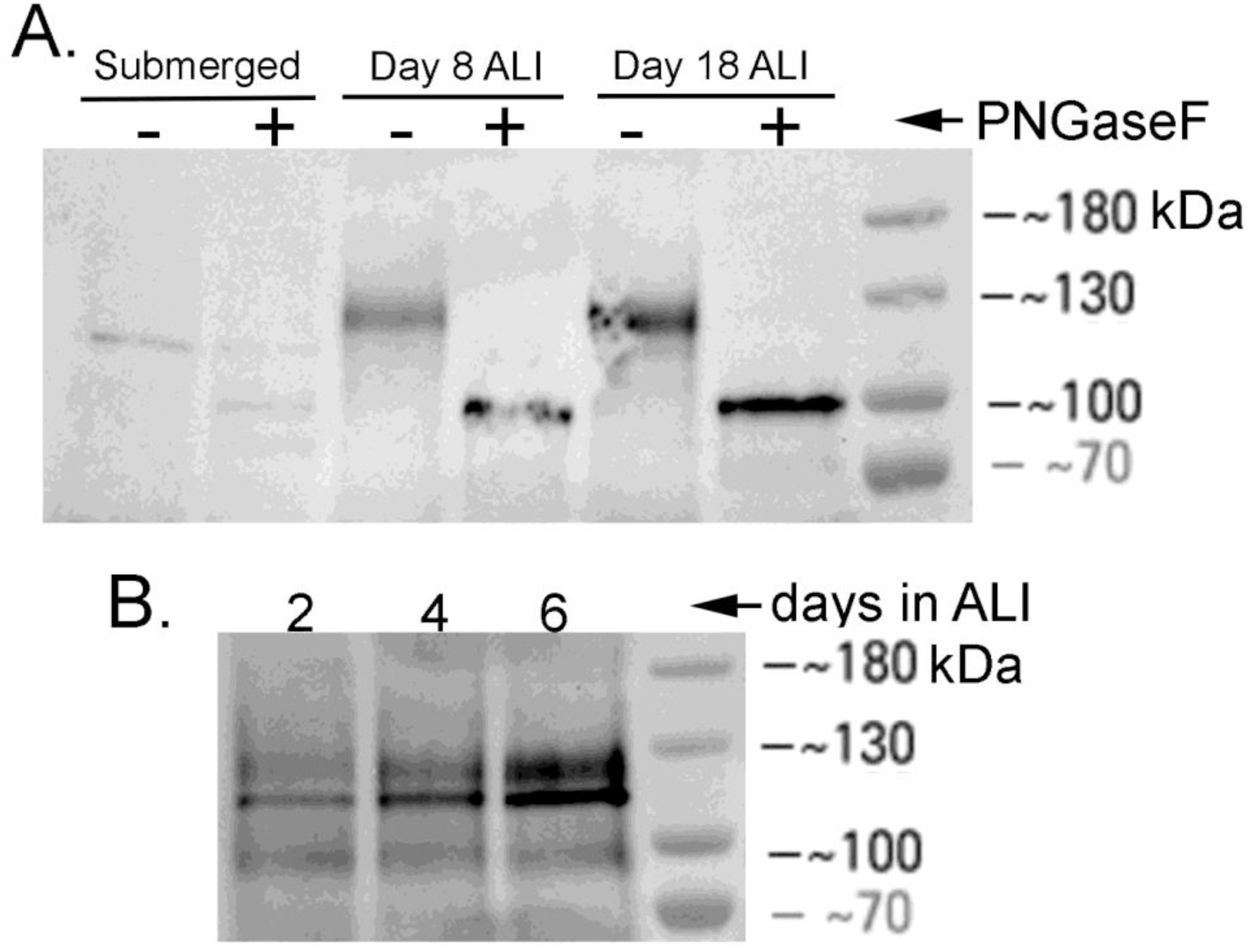
A.) Human nasal epithelial cells were grown under submerged (progenitor maintenance conditions) cell growth or subjected to air-liquid interface growth conditions. Cells were harvested at the indicated times and analyzed by immunoprecipitation followed by western blot analysis. As can be seen progenitor cells do not appear to express ACE2 protein and that the process of differentiation induces expression. PNGaseF treatment reveals that ACE2 exists as a glycoprotein. B.) To determine when ACE2 protein expression occurs during the differentiation process, HNECs were allowed to develop under ALI conditions. Cells were harvested at the indicated times and analyzed by immunoprecipitation and western blotting for ACE2 expression. As seen in the figure, ACE2 protein is apparent at day 2 and levels rise through days 4 and 6.

### ACE2 protein is both cytosolic and membrane-bound in ALI tissue

For our analysis of ACE2 protein we continued to use two different antibodies, one to the N-terminus (ProSci 3227) and the other to the C-terminus region (ProteinTech 66699). Although the two antibodies displayed the same band pattern when used in our western blot analyses they surprisingly exhibited distinct and overlapping patterns via fluorescent immunohistochemistry of ALI tissue (figure 5). Each antibody displayed signals in the apical cytosol of ciliated cells but only the ACE2 C-term antibody exhibited plasma membrane localization (figure 5). In basal cells we detected ACE2 localized to the basolateral region of the cytosol in these cells. It is important to note that SARS-CoV appears to be able to infect and be shed from the basolateral compartment [19]. It remains to be seen whether SARS-CoV-2 also productively infects the basal cells.

**Figure 5:**
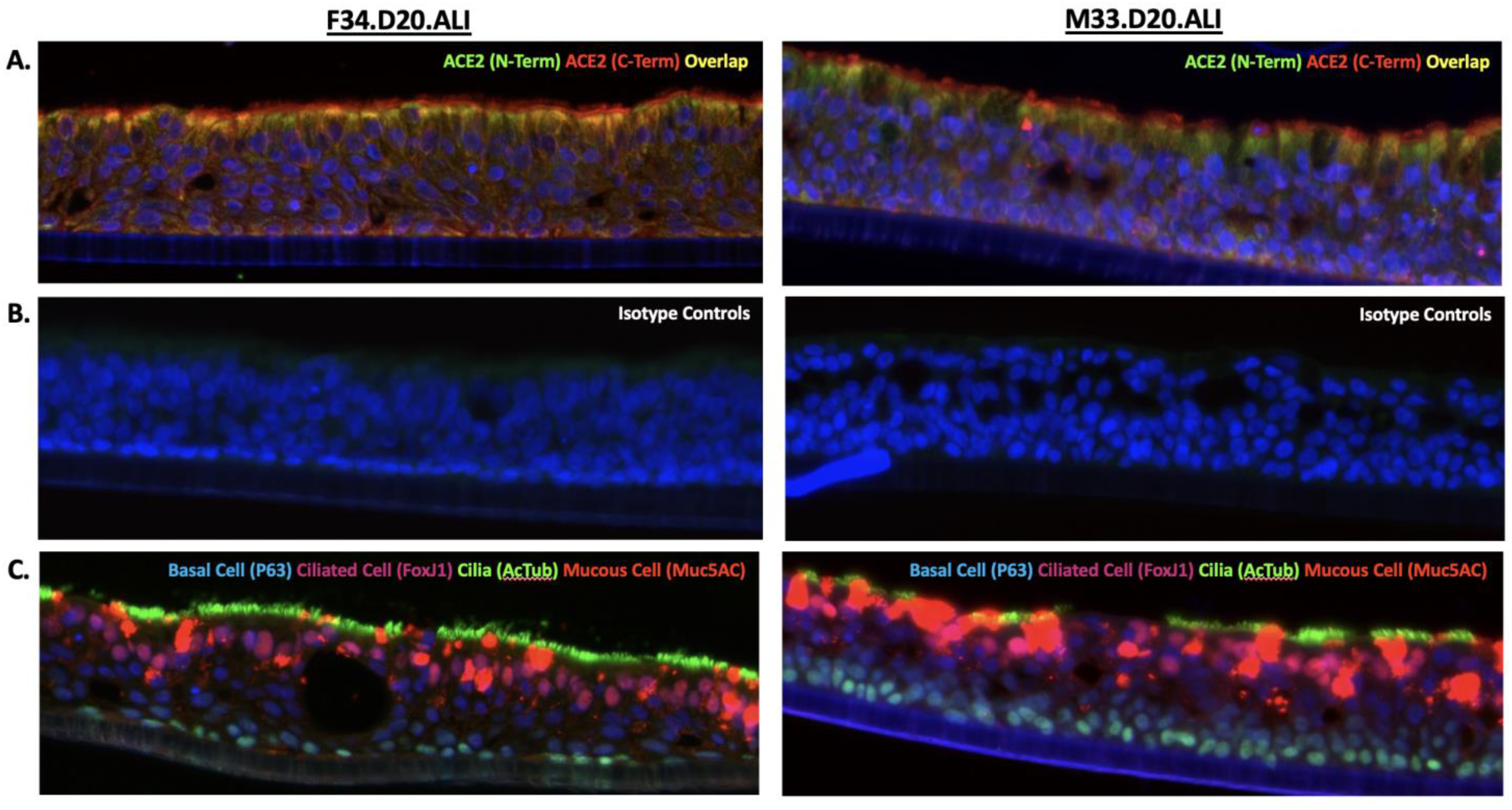
ACE2 protein localization was accomplished by fluorescent immunohistochemistry. We chose two different antibodies, one to the N-terminus and the other to the C-terminal half of the protein. Although both antibodies displayed the same signals when used in our western blot analysis they display distinct and overlapping signals via fluorescent immunohistochemistry. We observed ACE2 located both in the basal cell compartment and the ciliated cell compartment, but minimal signal in the central region of tissue. The apical cytosol of ciliated cells contained a mixture of individual signals and overlapping signals from both ACE2 antibodies. Only the ACE2 C-term antibody displayed plasma membrane signal on ciliated cells. When we compared ACE2 IHC patterns between ALI tissues derived from a 34-year old female to a 33-year old male we observed similar patterns but with varying intensity suggesting possible variation in the abundance of ACE2 protein in the airway between individuals. (A) ACE2 immunohistochemistry. (B) Rabbit and Mouse isotype controls (C) Immunohistochemical stain identifying progenitor cells (basal cells) and the differentiated cell types.

### ACE2 expression between donor ALI tissues

The membrane-bound and cytosolic localization of ACE2 signal was similar between donor ALI tissue derived from either a 34-year old female or a 33-year old male (figure 5). Although the localization pattern was similar we observed a reduction in the overall intensity of total ACE2 signal from the 33-year old male donor. These data suggest that the total amount of ACE2 protein in the airway may vary between individuals.

## DISCUSSION

Full-length ACE2 (variants 1,2) in addition to two predicted isoforms were cloned from well differentiated ALI tissues derived from two independent donors (F34, M33). This study is the first to identify and clone transcripts for the two predicted ACE2 isoforms vX1 and vX2. Whether or not these novel isoforms are specific to the airway is still to be determined. It is important to note that these two variants of full length ACE2 have completely unique residues at their extreme C-termini and are not mere truncations through exon deletion. Since each ACE2 isoform, including full-length, contain unique C-term protein motifs this indicates the possibility that each ACE2 isoform may possess a unique function.

To increase the rigor of our study we utilized two different ACE2 antibodies towards opposite regions of the protein (N-terminus and C-terminus) for our immunoblotting and immunofluorescent analyses. When single ACE2 isoforms were transfected into HEK293 cells and subsequent cell lysates were immunoblotted for ACE2, both antibodies gave identical signals, indicating that both antibody epitopes are present on every isoform. Immunofluorescence for ACE2 in transfected HEK293 cells revealed that full length ACE2 v1,2 localized to the plasma membrane, vX1 and vX2 also localized to the plasma membrane but at a reduced intensity with many less foci. In contrast to v1,2 both vX1 and vX2 showed cytosolic staining at a perinuclear location. Despite these studies being performed in immortalized cells rather than the primary airways cells that they were cloned from, the data undoubtedly demonstrates the intrinsic ability of these isoforms to localize to discrete areas of the cell.

An additional finding of these results was our immunofluorescent studies with ALI tissue which revealed that the two antibodies give distinct and overlapping patterns. Previous studies have shown that ACE2 protein localizes within the plasma membrane of ciliated cells [19]. We corroborated those observations and expanded on them by demonstrating ACE2 protein is also found within the cytosol of ciliated cells, as well as the cytosol of basal cells. Interestingly, we discovered plasma membrane ACE2 is only observed using the antibody towards the C-terminus region, suggesting that the epitope for the N-terminus antibody is blocked, possibly through an association with some unknown partner. Furthermore, the cytosol of ciliated cells exhibited green and yellow signals, corresponding to the N-terminus (Green) and an overlap of both N-term/C-term signals (Yellow). Altogether, the ciliated cells in ALI tissue displayed two areas of ACE2 localization with three distinct ACE2 signals (Cytosol/Green/N-term, Cytosol/Yellow/ N-term/C-term, Membrane/Red/C-term).

There are multiple scenarios that may explain the IHC patterns we observed in ALI tissue. One such explanation could be that the mixed signals are all the same ACE2 isoform in three stages of maturation or recycling and at each stage a distinct region of the ACE2 protein interacts with a specific partner. Another potential hypothesis is that these three signals represent the three individual isoforms functioning within their respective roles, and that the variation in N-term versus C-term signals are due to each isoform interacting with unique proteins. Pertaining to what the distinct roles of isoforms may be, previous studies demonstrated that soluble ACE2 maintains the ability to catabolize angiotensin II, suggesting that cytosolic, membrane bound or soluble ACE2 can potentially function in the resolution of inflammation in the airway [2]. Membrane bound ACE2 has been observed to function in cellular adhesion through interactions with integrins, indicating another potential role in airway tissue [11]. There is also the fascinating potential for undiscovered airway-specific substrates and functions of individual ACE2 isoforms.

The distinct ACE2 IHC patterns became more compelling upon the realization that the intensity of these signals seemed to vary between ALI tissues derived from different donors. We found that the ACE2 IHC patterns in ALI tissue isolated from our male donor with a history of smoking (M33) exhibited a great reduction in the overall intensity and amount of signal when compared to ALI tissue obtained from our female, nonsmoker donor (F34). Despite the small sample size of our study the implication stands that these parameters can vary among the population and perhaps the status of these airway ACE2 parameters play a direct role in COVID-19 susceptibility and severity.

We have detected that the amount of membrane bound and cytosolic ACE2 in ALI tissue varies between individual donor sources and we predict that production of soluble ACE2 will also vary. Previous studies demonstrated that soluble ACE2 interacts with the SARS-CoV spike protein and that increasing the concentration of soluble ACE2 conferred resistance to experimental SARS-CoV infection. The group also determined that ACE2 must be membrane-bound in order to facilitate entry of the SARS-CoV pathogen [16]. These studies support the postulate that having high concentrations of soluble ACE2 in the airway mucous layer will confer some level of resistance to SARS-CoV-2 infection, as each soluble ACE2 protein would in essence act as a ‘sponge’ for a viral particle. Expanding on this hypothesis, the consequence for an individual having high levels of membrane bound ACE2 is an increase in available entry points for viral particles and thereby an increase in susceptibility to SARS-CoV-2 infection. It’s conceivable that the wide variability in the severity of COVID-19 infections lie within the airway ACE2 status of the patients. Contributing to whether ACE2 is a functional target for SARS-CoV-2 include the potential for differential isoform expression, altered distribution in the cell between membrane-bound, cytosolic or the secreted form as well as differences in post-translational modification. It is in the interest of public health to understand how these ACE2 parameters affect the susceptibility of airway tissue to SARS-Cov-2 infection, in addition to the potential impact that diet, environment, genetics, prescription and over-the-counter medications may have on modulating these ACE2 parameters. Perhaps there are readily available, FDA-approved solutions that can help decrease susceptibility to COVID-19.

## METHODS

### HNEC Isolation and Expansion

Human nasal epithelia cells (HNECs) were isolated through brushing of inferior and middle nasal turbinates with interdental brushes (DenTek). Brushes were removed from the handle and placed into an enzymatic digestion buffer (ACF Enzymatic Dissociation Solution, StemCellTechnologies), briefly vortexed followed by incubation at 37C for 15 minutes. After digestion cells were pelleted through centrifugation at 200 x g for 5min, pellets were suspended in basal cell expansion media (PneumaCult Ex Plus Media, StemCellTechnologies) and incubated at 37C/5%CO_2_. Culture media was refreshed every two days, HNECs were collected and counted between 8-10 days of culture. HNECs were then either seeded directly onto transwell inserts for ALI differentiation or banked in liquid nitrogen for future experiments. Protocols for isolation of HNEC specimens from human subjects is approved by our Institutional Review Board and all donors provided informed consent.

### ALI Differentiation

HNECs were seeded onto semi-permeable polyester tissue culture-treated inserts that were 12mm in diameter and 0.4uM pore size (Corning) and cultured in expansion media until 100% confluence, confirmed by visualization of HNECs via phase contrast microscopy. Once confluent monolayer formed initiation of the air-liquid interface began by removing media from the transwell insert and replacing the expansion media in the lower-chamber with differentiation media (PneumaCult ALI Medium, StemCellTechnologies). Every 48 hours of ALI culture the media was refreshed and the transwell insert was rinsed with 500 µL PBS to prevent excessive mucous accumulation. Differentiation of ALI tissue was confirmed by visualization of beating cilia via phase contrast microscopy.

### ALI Tissue Fixation and Sectioning

ALI tissue was fixed in 10% buffered formalin at 4°C for 12-16 hours then dehydrated via 20-minute graded alcohol washes (70%-80%-95%-100%). Fixed and dehydrated ALI tissue was then embedded in paraffin wax blocks for sectioning via microtome.

### Hematoxylin and Eosin

Histological sections were deparaffinized in xylene and rehydrated via 5-minute graded alcohol washes (100%-95%-80%-70%). Sections were placed in hematoxylin for 1 minute, rinsed in tap water, dehydrated and stained with Eosin for 30 seconds. Lastly samples were cleared with xylene before applying coverslips. Staining was visualized and imaged via ECHO Revolve fluorescent microscope.

### Immunohistochemistry

Histological sections were deparaffinized in xylene and rehydrated via 5-minute graded alcohol washes (100%-95%-80%-70%-H_2_O). Antigen retrieval was performed in a digital pressure cooker (InstaPot) with slides submerged in desired antigen retrieval buffer. Sections were blocked with 5% BSA-TBST for one hour at room temperature. Primary antibodies were suspended at desired dilutions in 1% BSA-TBST and applied for one hour at room temperature, dilutions of individual antibodies can be found in table 2. Secondary antibodies conjugated to either 488 or 555 were applied at 1:1000 dilution in 1% BSA-TBST for one hour at room temperature. DAPI was applied and sections were cover-slipped and sealed. Immunofluorescence was visualized and imaged via an ECHO Revolve fluorescent microscope running the ECHO Pro imaging application.

**Table 1:**
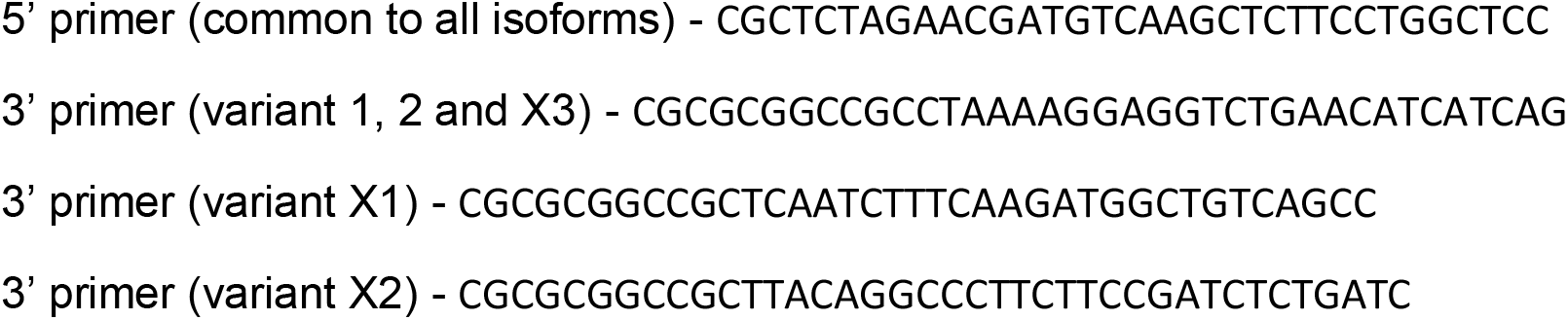
Primers used for the isolation of ACE2 variants.

**Table 2:**
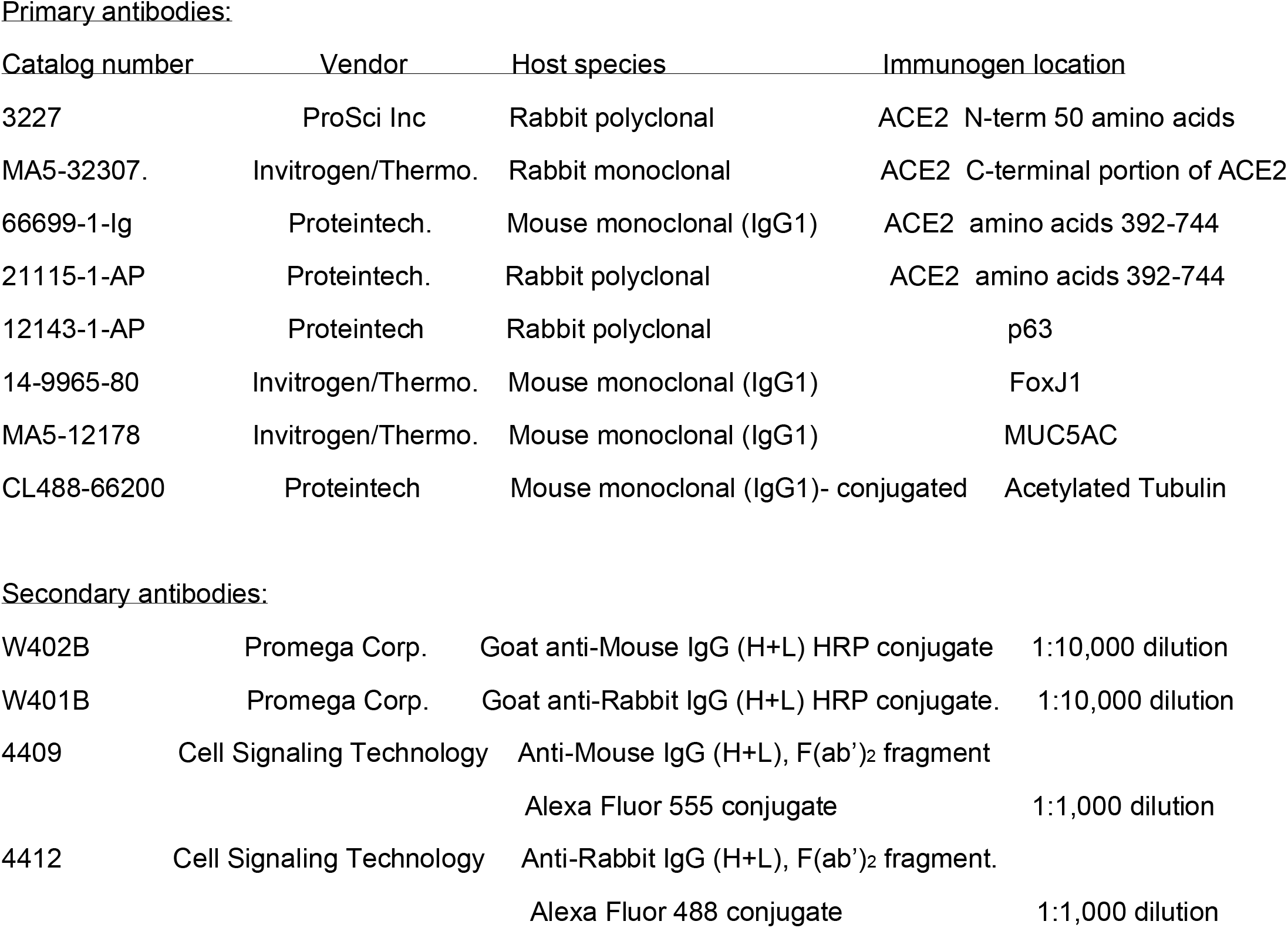
ACE2 Antibodies used in this study.

| Catalog number | Vendor | Host species | Immunogen location |
| --- | --- | --- | --- |
| 3227 | ProSci Inc | Rabbit polyclonal | ACE2 N-term 50 amino acids |
| MA5-32307. | Invitrogen/Thermo. | Rabbit monoclonal | ACE2 C-terminal portion of ACE2 |
| 66699-1-Ig | Proteintech. | Mouse monoclonal (IgG1) | ACE2 amino acids 392-744 |
| 21115-1-AP | Proteintech. | Rabbit polyclonal | ACE2 amino acids 392-744 |
| 12143-1-AP | Proteintech | Rabbit polyclonal | p63 |
| 14-9965-80 | Invitrogen/Thermo. | Mouse monoclonal (IgG1) | FoxJ1 |
| MA5-12178 | Invitrogen/Thermo. | Mouse monoclonal (IgG1) | MUC5AC |
| CL488-66200 | Proteintech | Mouse monoclonal (IgG1)- conjugated | Acetylated Tubulin |
| Secondary antibodies: |  |  |  |
| W402B | Promega Corp. | Goat anti-Mouse IgG (H+L) HRP conjugate | 1:10,000 dilution |
| W401B | Promega Corp. | Goat anti-Rabbit IgG (H+L) HRP conjugate. | 1:10,000 dilution |
| 4409 | Cell Signaling Technology | Anti-Mouse IgG (H+L), F(ab')<sub>2</sub> fragment |  |
|  |  | Alexa Fluor 555 conjugate | 1:1,000 dilution |
| 4412 | Cell Signaling Technology | Anti-Rabbit IgG (H+L), F(ab')<sub>2</sub> fragment. |  |
|  |  | Alexa Fluor 488 conjugate | 1:1,000 dilution |

### ACE2 Protein Analysis

*Protein extraction for total cell extracts:* Cells were washed and harvested in PBS. The cell pellet was frozen at −70°C until use. Cells were resuspended in total cell extract/immunoprecipitation buffer (TCE/IP buffer; 50 mM Tris-Cl pH 7.5 @ 4°C, 150 mM NaCl, 1 mM EDTA, 1% Triton X-100, protease inhibitors added immediately before use) and incubated on ice for 15 min. Cells were disrupted in a Bioruptor (diagenode, UCD 200). Settings were on high, 30 sec. on and 30 sec. off for 5 min. Insoluble debris was pelleted and the supernatant (TCE) used for further analysis.

#### Immunoprecipitation protocol

500 µg of TCE protein was added to 1 ml of TCE/IP buffer. 1µg of antibody (Proteintech, 21115-1-AP) was added and the solution was incubated overnight at 4°C on a rotating platform. 20 µl of a Protein A sepharose (nProtein A Sepharose 4 fast flow, GE Healthcare, 17-5280-01) suspension was added and incubated for 1 hour at 4°C on a rotator. The protein A suspension was pelleted and washed three times for 5 minutes each with TCE/IP buffer. For deglycosylation studies the pellets were resuspended in 200 µl of TCE/IP buffer, divided into two equal volumes and pelleted. The pellets were treated according to protocol and incubated for 1.5 hours at 37°C plus or minus PNGaseF. Protein A pellets were resuspended in SDS-PAGE buffer and subjected to gel electrophoresis.

#### SDS-PAGE electrophoresis

8%-16% precast Novex Tris-Glycine gels (ThermoFisher Scientific,Waltham, MA) were used following standard protocols established in our laboratory [20].

#### Western blot analysis

Electrophoretically separated protein was transferred to PVDF. Standard protocols were used to perform the western blot analysis [21]. ECL chemiluminescent western blotting detection system (Amersham Corp., Buckinghamshire, UK) was used to develop positive signals. Signal acquisition was accomplished using a ThermoFisher Ibright fl1500 imaging system. Immunostaining antibody (Invitrogen; MA5-32307.

#### Deglycosylation of ACE2 protein

PNGase F, which is an enzyme effective in the removal of almost all N-linked oligosaccharides from glycoproteins was purchased from New England Biolabs. Vendor protocols were followed for the use of PNGase F.

### Cloning and Transfection studies

cDNA libraries were constructed from RNA derived from HNECs grown either in submerged conditions in PneumaCult Ex Plus media or after ALI differentiation from samples taken between 15 to 25 days. RNA was isolated from cells using the NucleoSpin RNA Plus kit of Clontech Laboratories (Takara Bio Company). cDNA was constructed using the In-Fusion SMARTer Directional cDNA Library Construction kit of Takara Bio. PCR was then used to isolate clones of ACE2 using primers listed in table 1. A common 5’ primer encoding an XbaI restriction site was used along with isoform specific 3’ primers containing a NotI restriction enzyme site. PCR amplification was accomplished with Q5 High-Fidelity DNA Polymerase from New England Biolabs. PCR products derived from the cDNA libraries were digested with the indicated enzymes and ligated into the pCINeo vector of Promega Corporation. *E. coli* (New England Biolabs, DH5alpha) were transformed with the ligation reactions. Individual clones were identified by colony PCR using primers within the pCINeo vector (figure 1A). Verification of DNA sequence was accomplished by Genewiz (South Plainfield, NJ). HEK-293 cells (ATCC CRL-1573) were used for transfection studies. Identified and characterized plasmids containing the ACE2 isoform open reading frames were transfected into the HEK-293 cells using TurboFect reagent (ThermoFisher Corp.). 24 hours post-transfection the cells were harvested and processed for analysis by western blot according to established procedures [21]. For fluorescence microscopy, 24-hour transfected cells were trypsinized and replated on coverslips. 24 hours later the coverslips were processed for visualization. (Antibodies used; ProSci 3227, Proteintech 66699).

